# Peroxisomal β-oxidation enzyme, DECR2, regulates lipid metabolism and promotes treatment resistance in advanced prostate cancer

**DOI:** 10.1101/2022.11.05.515262

**Authors:** Chui Yan Mah, An Dieu Trang Nguyen, Takuto Niijima, Madison Helm, Jonas Dehairs, Feargal J Ryan, Natalie Ryan, Ian G Mills, Johannes V Swinnen, David J Lynn, Zeyad D Nassar, Lisa M Butler

**Affiliations:** South Australian Immunogenomics Cancer Institute and Freemasons Centre for Male Health and Wellbeing, University of Adelaide, Adelaide, Australia; Precision Medicine Theme, South Australian Health and Medical Research Institute, Adelaide, Australia; Department of Oncology, Laboratory of Lipid Metabolism and Cancer, KU Leuven, Leuven, Belgium; Patrick G Johnston Centre for Cancer Research, Queen’s University, Belfast, United Kingdom; Flinders Health and Medical Research Institute, Flinders University, Bedford Park, Australia

## Abstract

Peroxisomes are central metabolic organelles that have key roles in fatty acid homeostasis, including β-oxidation, and emerging evidence has linked aberrant peroxisome metabolism to cancer development and progression. While targeting mitochondrial β-oxidation in prostate cancer (PCa) has gained significant attention in recent years, the contribution of peroxisomal β-oxidation (perFAO) to PCa tumorigenesis is comparatively unexplored. Herein, we explored the therapeutic efficacy of targeting perFAO in PCa cells and clinical prostate tumours, and subsequently identified peroxisomal 2,4-dienoyl CoA reductase 2 (DECR2), as a key therapeutic target. DECR2 is markedly upregulated in clinical PCa, most notably in metastatic castrate-resistant PCa. Depletion of DECR2 significantly suppressed proliferation, migration, and 3D growth of a range of CRPC and enzalutamide-resistant PCa cell lines, and inhibited LNCaP tumour growth and proliferation *in vivo*. Using transcriptomic and lipidomic analyses, we determined that DECR2 influences cell cycle progression and lipid metabolism to enable tumour cell proliferation. We further demonstrated a novel role for perFAO in driving resistance to standard-of-care androgen receptor pathway inhibition, using genetic and pharmacological approaches to alter DECR2/perFAO in treatment-resistant PCa cells. Our findings highlight a need to focus on peroxisomes to suppress tumour cell proliferation and reveal new therapeutic targets for advanced, treatment-resistant PCa.

## INTRODUCTION

Prostate cancer (PCa) remains the most diagnosed malignancy and the leading cause of cancer-related deaths in men globally^1^. One of the main hurdles for the treatment of PCa is overcoming resistance to current androgen-targeting agents, which form the mainstay of therapy for locally advanced and metastatic PCa. Despite the development of potent androgen receptor (AR) pathway inhibitors, including enzalutamide (ENZ), these agents are not curative and patients with castrate-resistant prostate cancer (CRPC) eventually succumb to this disease. Targeting cancer metabolism has gained increasing attention as an attractive strategy to overcome resistance to AR-targeted therapies^2–7^.

Altered lipid metabolism is a well-characterised hallmark of PCa and, accordingly, significant research efforts have been made to target *de novo* lipogenesis and lipid uptake pathways^7,8^. Now, there is a growing body of evidence that implicates fatty acid oxidation (FAO, or β-oxidation) as a critical aspect of lipid metabolism that drives PCa progression and treatment resistance^9–12^, irrespective of fatty acid source. Despite the complexity of the FAO pathway, most drug development approaches have focused entirely on targeting mitochondrial carnitine palmitoyltransferase I (CPT1), the rate-limiting enzyme of mitochondrial β-oxidation^13–15^. Indeed, our previous work demonstrated therapeutic efficacy in targeting mitochondrial β-oxidation using a pharmacological agent, etomoxir (CPT1 inhibitor) in our established patient-derived explant (PDE) model^12^. With this observation, we also uncovered a novel target of mitochondrial β-oxidation, DECR1, the rate-limiting enzyme of an auxiliary pathway of polyunsaturated fatty acid (PUFA) oxidation^12^. On the basis of this discovery, we were motivated to investigate and characterise its peroxisomal counterpart, peroxisomal 2,4-dienoyl-CoA reductase 2 (DECR2), and its role in PCa.

Peroxisomes are organelles that regulate the synthesis and turnover of complex lipids, including the β-oxidation of very-long chain fatty acids (VLCFA) (**Figure 1a**), synthesis of bile acids and ether lipids (such as plasmalogens), α-oxidation of branched-chain fatty acids (BCFA), and regulate cholesterol biosynthesis. Although both the peroxisome and mitochondria have similar functions (for example, both organelles can degrade fatty acids and produce/scavenge reactive oxygen species), it is becoming increasingly clear that peroxisomes are indispensable organelles that are essential for cellular well-being. For instance, peroxisomes are the sole organelles in humans able to break down VLCFAs and the only ones performing α-oxidation. Despite the critical roles of peroxisomes in lipid metabolism, the functional effects of peroxisomal β-oxidation (perFAO) in cancer are not well recognised and not as intensively studied as those of mitochondria. To this end, the most well-characterised perFAO enzyme in PCa is α-methylacyl-CoA racemase (AMACR; involved in β-oxidation of BCFA). AMACR is consistently overexpressed in PCa and is associated with increased PCa risk^16^. More importantly, AMACR is highly specific for PCa and thus has been exploited as a PCa-specific biomarker^17^. In a more recent study, Itkonen et al. reported that peroxisomal enoyl-CoA Delta Isomerase 2 (ECI2; involved in β-oxidation of unsaturated fatty acids) was significantly upregulated in human PCa and is associated with poor overall patient survival^18^. Herein, we demonstrate the therapeutic efficacy of targeting perFAO *in vitro* and in our patient-derived prostate tumour explants (PDE) to provide the first clinically relevant evidence for targeting perFAO in PCa. We subsequently identified DECR2 as robustly overexpressed in advanced and metastatic PCa tissues and uncovered its function as a regulator of cell cycle progression and lipid metabolism. Finally, we provide evidence for a role of DECR2 and perFAO in treatment resistance, indicating a novel therapeutic vulnerability for CRPC.

**Figure 1.**
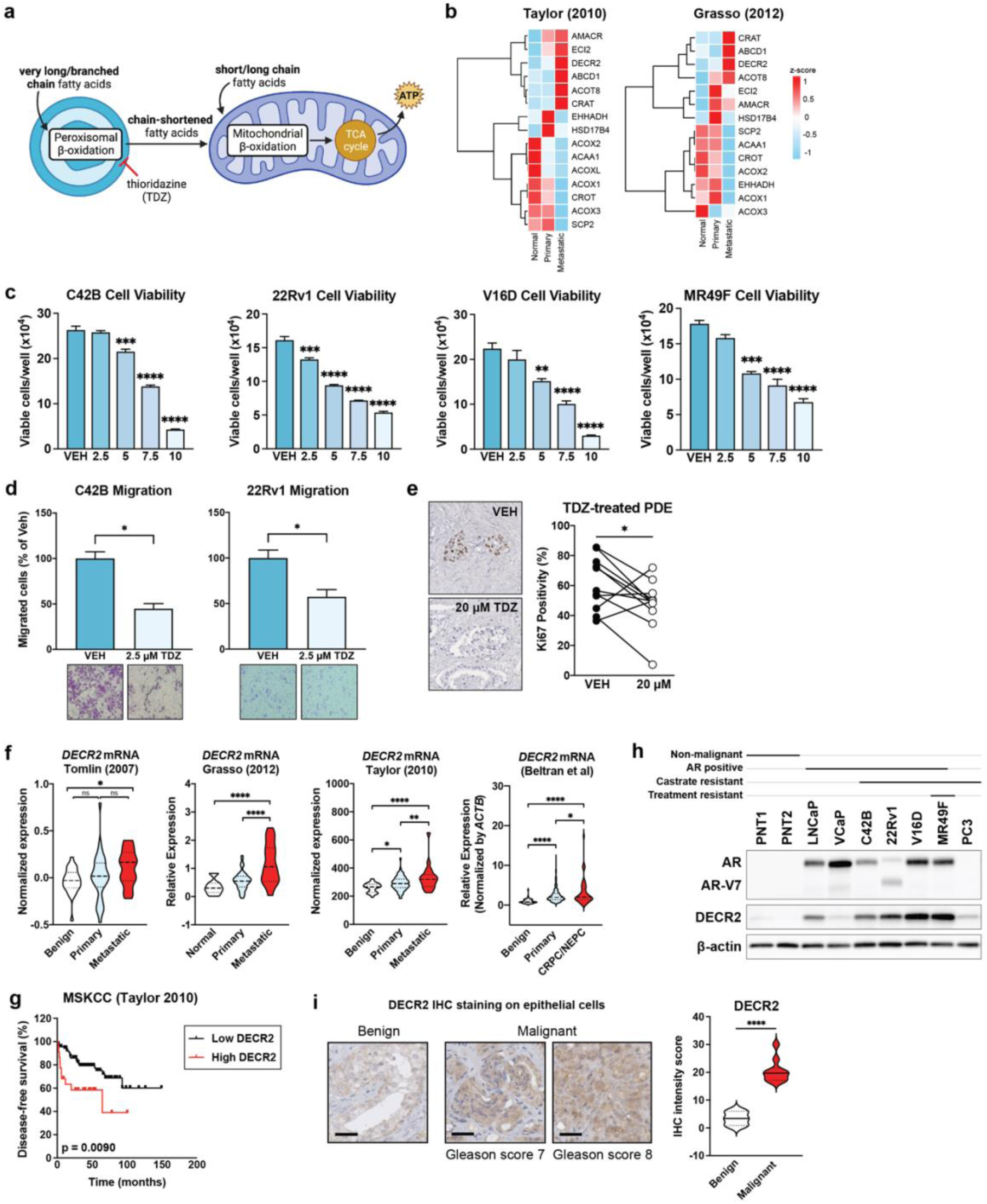
DECR2 is overexpressed in prostate cancer. **(a)** Illustration of fatty acid oxidation in the peroxisome and mitochondria. Thioridazine (TDZ) is an inhibitor of perFAO. **(b)** Heatmap of peroxisomal β-oxidation (perFAO) gene expression in Taylor and Grasso cohorts. We manually curated a list of perFAO genes based on Gene Ontology pathway. Cell viability of **(c)** castrate-resistant C42B, 22Rv1 and V16D, and enzalutamide-resistant MR49F prostate cancer cell lines across a range of TDZ doses. **(d)** C42B and 22Rv1 prostate cancer cell lines treated with 2.5 µM TDZ were assessed for cell migration using transwell migration assay. **(e)** Immunostaining for proliferative marker Ki67 in vehicle (VEH) or TDZ-treated (20 μM) patient-derived explants (PDEs). Immunohistochemical staining and quantification of the proliferative marker Ki67 is shown (*n* = 11). **(f)** DECR2 expression with respect to tumour progression in four independent datasets. DECR2 levels were analysed in normal, primary, and metastatic castrate-resistant or neuroendocrine tissue samples. **(g)** The association of DECR2 expression and disease-free survival in the MSKCC (Taylor) cohort. **(h)** DECR2 protein expression in non-malignant prostate cell lines (PNT1 and PNT2) and prostate cancer cell lines (LNCaP, VCaP, C42B, 22Rv1, V16D, PC3), including enzalutamide-resistant prostate cancer cell line (MR49F). **(i)** Left: Representative DECR2 IHC staining of benign prostate tissues and prostate cancer tissues. Scale bar, 50 µm. Right: DECR2 protein expression in a validation cohort consisting of benign prostate tissues (*n* = 3) and prostate cancer tissues (*n* = 10). All cell line data are representative of at least 2 independent experiments and presented as mean ± s.e.m of triplicate wells. Statistical analysis was performed using ordinary one-way ANOVA or two-tailed student’s t-test. Data in (g) were statistically analysed using a two-sided log-rank test. \**p* < 0.05, \*\**p* < 0.01, \*\*\**p* < 0.001 and \*\*\*\**p* < 0.0001.

## RESULTS

### Peroxisomal β-oxidation enzyme, DECR2, is overexpressed in prostate cancer

Little is known about targeting the enzymes of perFAO or their expression in PCa. Accordingly, we evaluated the expression of a set of peroxisome-related genes (obtained from the KEGG database) in the Taylor cohort^19^ composed of primary (*n* = 131) and metastatic (*n* = 19) tumour tissues and noticed a significant variation in the expression of these genes with tumour progression (**Supplementary Figure 1a**). Next, we sought to identify genes that are involved in perFAO. To achieve this, we investigated GO terms (Molecular Signatures Database, MSigDB) for perFAO and manually curated a list of *n* = 15 genes that were involved in perFAO, and examined their expression in the Taylor and Grasso^20^ (primary *n* = 59, metastatic CRPC *n* = 35) cohorts (**Figure 1b**). Although we observed variability in their expression, this revealed several perFAO genes: *ECI2, DECR2, ABCD1, CRAT,* and *ACOT8* (**Figure 1b**) involved in regulation of fatty acid metabolism and energy homeostasis that were consistently upregulated in both cohorts in metastatic tissues, suggesting an important role for perFAO in PCa.

In light of this, we evaluated the efficacy of targeting perFAO using a potential clinical candidate agent and inhibitor of perFAO, thioridazine (TDZ)^21,22^, in CRPC and treatment-resistant PCa cells (**Figure 1a**). TDZ is a first-generation antipsychotic drug that was withdrawn from the global market in 2005 due to a well-defined risk of cardiac arrythmias. Despite that, TDZ continues to be used off-label for patients with severe or chronic schizophrenia who are refractory to other treatment options^23^. In recent years, TDZ has been increasingly used as a perFAO inhibitor^22,24^, likely through its inhibitory effects on Cytochrome P450 enzymes^25,26^. TDZ induced a dose-dependent reduction in cell viability of CRPC cells (C42B, 22Rv1, V16D cell lines) and acquired ENZ-resistant MR49F cells (**Figure 1c**). Furthermore, we tested the ability of TDZ to impede cell migration of C42B and 22Rv1 CRPC cells, and showed significantly reduced migration at a low dose of 2.5 µM of TDZ (**Figure 1d**).

TDZ significantly and dose-dependently decreased colony formation ability, and induced apoptosis and cell death in CRPC C42B, V16D and ENZ-resistant MR49F cell lines (**Supplementary Figure 1b,c**). Our recent report demonstrated the efficacy of targeting mitochondrial β-oxidation using the chemical inhibitor, etomoxir^12^, using our well-defined patient-derived explant (PDE) model that recapitulates the complexity of the clinical tumour microenvironment^27^. Herein, we evaluated the clinical efficacy of targeting perFAO using TDZ in PDE tissues and observed an overall significant reduction in cell proliferation by an average of 21.4 ± 4.0% (*n* = 11; *p* < 0.05), with only 2 patients showing no response (**Figure 1e**).

Our results demonstrated that TDZ is efficacious *in vitro* and *ex vivo* and provide proof-of-concept that targeting perFAO may be a promising therapeutic strategy. However, no specific inhibitors of perFAO currently exist. In view of our recent discovery of the role of mitochondrial DECR1 in PCa^12^ and its upregulation in metastatic tissues, we focused our attention on DECR2 as a key enzyme involved in perFAO. We further validated the overexpression of DECR2 mRNA in the Tomlin^28^ (*n* = 51 primary and metastatic PCa) and Beltran^29^ (CRPC *n* = 34, neuroendocrine PCa *n* = 15) cohorts and observed significantly higher levels of DECR2 in metastatic tissue compared with primary (Beltran cohort) or normal tissue (Tomlin and Beltran cohorts) (**Figure 1f**). In line with this observation, DECR2 gene copy number gain was evident in several clinical PCa datasets (acquired from cBioportal; **Supplementary Figure 1d**). Higher DECR2 levels were also significantly associated with biochemical recurrence in the MSKCC (metastatic CRPC) cohort (**Figure 1g**). Next, we examined DECR2 protein expression in a panel of cell lines: DECR2 levels were low in AR-positive, androgen-dependent LNCaP cells and AR-positive CRPC C42B cells, intermediate in AR-positive CRPC 22Rv1 cells, and high in AR-positive CRPC V16D and ENZ-resistant MR49F cell lines (**Figure 1h**). Similarly to publicly available datasets, we also observed an increase in DECR2 expression in malignant PCa tissues (*n* = 10) compared with benign tissues (*n* = 3), as assessed using quantitative immunohistochemistry staining analysis (**Figure 1i**). Consistent with its known function, we confirmed DECR2 localisation in the peroxisome using immunocytochemistry (**Supplementary Figure 1e**). Furthermore, we provide evidence for the perFAO selectivity of TDZ by demonstrating that overexpression of DECR2 markedly increased susceptibility of LNCaP cells to TDZ treatment, while TDZ had no effect on DECR2 knockdown cells (**Supplementary Figure 1f, g**).

### DECR2 targeting inhibits prostate cancer oncogenesis

The upregulation of DECR2 levels in metastatic CRPC compared to benign tissues suggests an important role for DECR2 in PCa growth and progression. Indeed, transient knockdown of DECR2 significantly suppressed viability and induced cell death in androgen-dependent LNCaP, CRPC 22Rv1 and V16D, and enzalutamide-resistant MR49F PCa cell lines (**Figure 2a**, **Supplementary Figure 2a**). No effect on cell viability was observed in non-malignant PNT1 prostate cells (**Supplementary Figure 2b**). Similarly, doxycycline (dox)-inducible knockdown of DECR2 with short hairpin RNA (shDECR2+Dox) significantly attenuated viability in LNCaP PCa cell lines (**Figure 2b**). In contrast, constitutive ectopic overexpression of DECR2 (hDECR2) in LNCaP cells significantly enhanced viability compared with vector control cells (hControl; **Figure 2c**). Additionally, dox-inducible knockdown of DECR2 markedly decreased LNCaP colony formation while stable overexpression of DECR2 increased colony formation (**Figure 2d, e**). Likewise, DECR2 knockdown significantly reduced migration of CRPC C42B and 22Rv1 PCa cells (**Supplementary Figure 2c**), while stable overexpression of DECR2 increased migration in LNCaP cells (**Figure 2f**). Finally, dox-induced shDECR2 (shDECR2+Dox) cells showed significantly reduced capacity for tumour growth and a trend towards reduced lung metastasis compared with non-dox-induced (shDECR2-Dox) cells using LNCaP orthotopic xenografts (**Figure 2g**; **Supplementary Figure 2d**). Inspection of the tumours also revealed significantly reduced cellular proliferation in shDECR2+Dox cells compared with shDECR2-Dox cells (**Figure 2h**). In contrast, overexpression of DECR2 in LNCaP cells significantly increased tumour growth compared with control cells (**Figure 2i**). In addition, analysis of detectable tumours from hDECR2 (*n* = 6) and hControl (*n* = 10) mice revealed a significant increase in tumour weight and lung metastasis of hDECR2 cells compared with control cells (**Figure 2i, j**).

**Figure 2.**
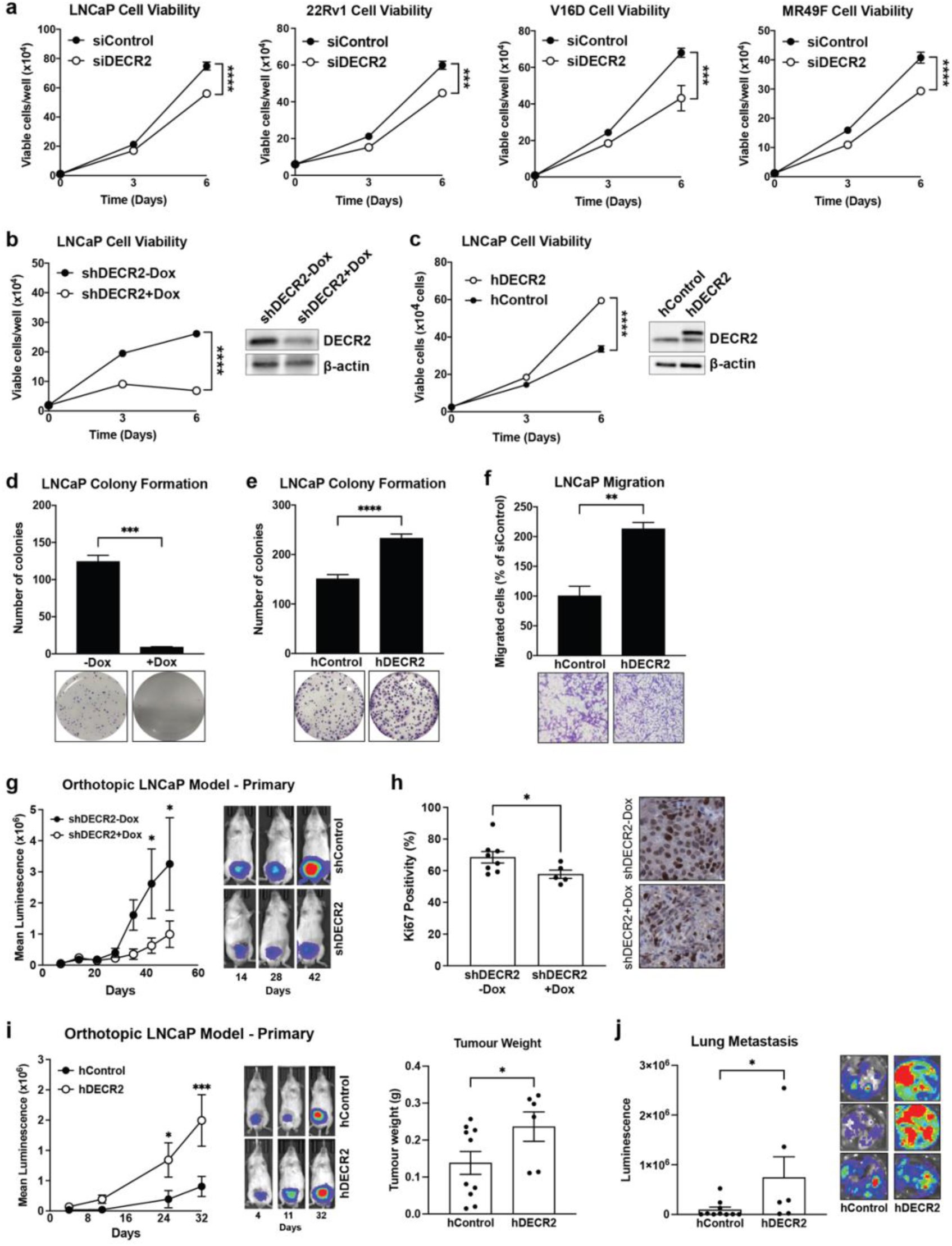
DECR2 knockdown inhibits prostate cancer cell growth in vitro and in vivo. **(a)** Cell viability of androgen-dependent LNCaP, castrate-resistant 22Rv1 and V16D, and enzalutamide-resistant MR49F prostate cancer cell lines subjected to siRNA-mediated DECR2 knockdown. **(b)** Cell viability of LNCaP cells with stable/inducible shRNA DECR2 knockdown (shDECR2) and **(c)** stable overexpression of DECR2 (hDECR2). **(d)** LNCaP colony formation was evaluated in cells with stable/inducible shRNA DECR2 knockdown (shDECR2) or **(e)** stable DECR2 overexpression (hDECR2). **(f)** LNCaP stable DECR2 overexpression (hDECR2) cell lines were assessed for cell migration using a transwell migration assay. **(g)** LNCaP cells with stable/inducible shRNA knockdown of DECR2 (shDECR2, *n* = 11) or control (shControl, *n* = 10) were analysed for orthotopic LNCaP tumour growth in mice, representative bioluminescent tumour images (right). **(h)** Ki67 quantification (left) and representative IHC staining (right) of orthotopic LNCaP tumours. Scale bar, 100 µm. This panel includes data from mice with sufficient sized tumours for analysis (shDECR2 *n* = 5, shControl *n* = 8). **(i)** Tumour growth and tumour weight of intraprostatically injected LNCaP cells with stable DECR2 overexpression (hDECR2, *n* = 6) or control (hControl, *n* = 10), representative bioluminescent tumour images (right). **(j)** Lung luminescence readings of stable DECR2 overexpression tumours in mice. All *in vitro* data are representative of at least 2 independent experiments and presented as mean ± s.e.m of triplicate wells. Statistical analysis was performed using ordinary one-way ANOVA or two-tailed student’s t-test: \**p* < 0.05, \*\**p* < 0.01, \*\*\**p* < 0.001 and \*\*\*\**p* < 0.0001.

### Depletion of DECR2 induces cell cycle arrest

To investigate the mechanism by which PCa cell growth and proliferation was attenuated by knockdown of DECR2, we carried out genome-wide transcriptional profiling of V16D and MR49F PCa cells (*n* = 6 biological replicates) subjected to a pooled siRNA-mediated knockdown of DECR2 (**Figure 3a**). Differential expression analysis identified > 8000 genes that were significantly (FDR < 0.05) differentially expressed in DECR2 knockdown cells compared to control cells (**Supplementary Data 1**). Gene Set Enrichment Analysis (GSEA) revealed a strong enrichment for GO-terms/pathways (MSigDB) related to the cell cycle and DNA replication and repair processes among downregulated genes in DECR2 knockdown cells compared with control cells (**Supplementary Data 1**). Other metabolic processes such as carboxylic acid pathways, branched-chain amino acid pathways and isoprenoid metabolic processes were enriched among genes upregulated in DECR2 knockdown cells (**Figure 3b**; **Supplementary Figure 3a; Supplementary Data 1**). GSEA analysis of the Hallmark pathway terms (MSigDB) identified E2F targets to be enriched among downregulated genes (**Figure 3c**). Accordingly, we examined the effect of DECR2 knockdown on cell cycle profile by flow cytometry. Knockdown of DECR2 induced cell cycle arrest at the G1/S phase in V16D and MR49F PCa cell lines (**Figure 3d**). Further, we validated our observations via qPCR in inducible DECR2 knockdown and overexpression cells (**Supplementary Figure 3b, c**). Stable overexpression of DECR2 in LNCaP cells showed the opposite effect where there were enhanced proportions of cells in S phase, consistent with an increase in cell proliferation (**Supplementary Figure 3d**). Likewise, TDZ induced G1/S phase cell cycle arrest dose-dependently in V16D and MR49F cells, particularly at the 10 µM dose (**Figure 3e**). In light of these findings, we assessed the effect of DECR2 knockdown on several cell-cycle related proteins/regulators in V16D and MR49F cells. We observed an increase in cyclin-dependent kinase inhibitors p21 and p27, and a decrease in cyclin-dependent kinase CDK4 (**Figure 3f**). Notably, we observed a decrease in phosphorylated retinoblastoma (pRb), a tumour suppressor protein in DECR2 knockdown cells compared to control cells (**Figure 3f**). We next evaluated whether the cyclin-dependent kinase (CDK) 4/6 inhibitor, ribociclib (Rib) could further enhance the effect of DECR2 knockdown. Indeed, Rib further abrogated viability of DECR2 depleted V16D and MR49F cell lines (**Figure 3g**). In contrast, stable overexpression of DECR2 in LNCaP cells rendered the cells more resistant to Rib compared to control cells (**Supplementary Figure 3d, e**). Finally, we examined the effect of perFAO inhibition via TDZ in combination with Rib on PCa cell viability. We found that TDZ further abrogated viability of V16D and MR49F cell lines when treated in combination with Rib compared to vehicle-treated cells or Rib alone (**Figure 3h**).

**Figure 3.**
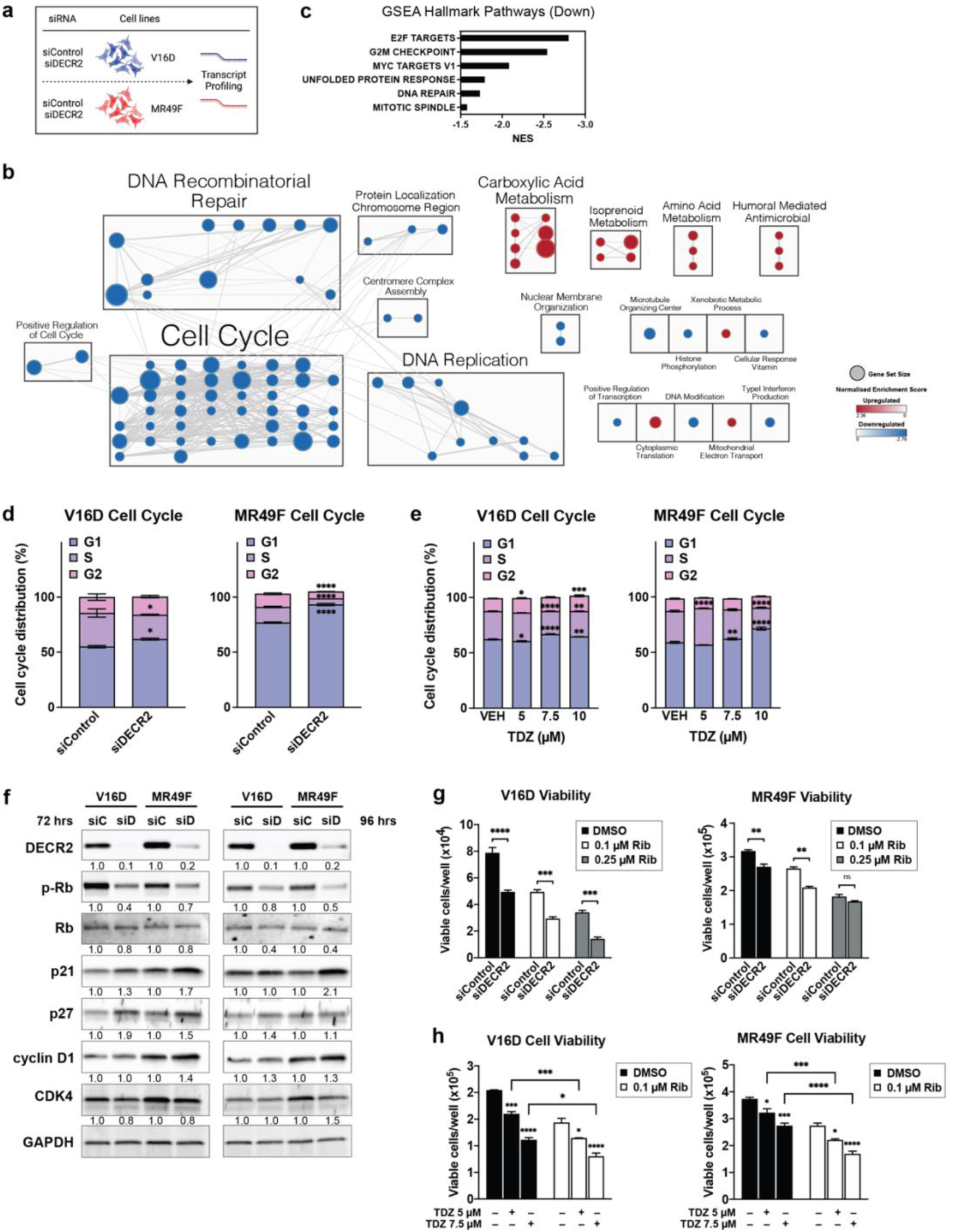
Transcriptomic analysis of the molecular mechanism of DECR2 function. **(a)** Schematic for DECR2-dependent RNA-seq-based changes in gene expression after DECR2 siRNA knockdown. **(b)** Gene interaction network of GSEA GO-terms (sourced from MSigDB, FDR < 0.01) enriched in up- (red) or down- (blue) regulated genes in DECR2 knockdown V16D and MR49F prostate cancer cells. Nodes represent gene sets and node size represents the number of genes in the gene set. Edges represent overlap between gene sets and edge width represents the number of genes that overlap (see **Supplementary Data 1**). **(c)** Bar chart of enriched GSEA Hallmark terms among downregulated genes in DECR2 knockdown cells. V16D and MR49F cell cycle distribution 96 h after **(d)** siRNA-mediated DECR2 knockdown **(e)** TDZ treatment. Data presented as percentage of cells in G1, S or G2 phase per sample. **(f)** Western blot analysis of a panel cell cycle-related protein markers in V16D and MR49F cells 72 and 96 h after DECR2 knockdown. GAPDH was used as loading control. Cell viability of **(g)** V16D and MR49F prostate cancer cells after DECR2 knockdown. **(h)** Cell viability of V16D and MR49F prostate cancer cells treated with TDZ (5 μM and 7.5 μM) and/or in combination with Rib (0.1 μM). All *in vitro* data are representative of at least 2 independent experiments and presented as mean ± s.e.m of triplicate wells. Statistical analysis was performed using ordinary one-way or two-way ANOVA: ns = non-significant, \**p* < 0.05, \*\**p* < 0.01, \*\*\**p* < 0.001 and \*\*\*\**p* < 0.0001.

One of the most strongly enriched pathways from our RNAseq data was lipid metabolism and fatty acid metabolic processes (**Figure 3b**; **Supplementary Figure 3a**). Consistent with these data, transcription factor enrichment analysis^30^ identified several lipid metabolism-related transcription factors (i.e., HNF4G, HNF4A, PPARG) that were significantly enriched among the top upregulated differentially expressed genes in DECR2 knockdown cells (**Supplementary Figure 3f**).

### DECR2 knockdown dysregulates lipid metabolism of prostate cancer cells

Given the biological role of DECR2 in perFAO, we further explored the impact of perturbing DECR2 on lipid metabolism. Knockdown of DECR2 in V16D and MR49F PCa cells for 4 days significantly induced neutral lipid accumulation, suggesting storage of lipids in lipid droplets (**Supplementary Figure 4a**). Likewise, TDZ dose-dependently increased neutral lipid accumulation in V16D and MR49F PCa cell lines (**Supplementary Figure 4b**). To understand how cellular lipid composition is altered by DECR2 knockdown, we carried out a global lipidomic analysis of LNCaP, V16D and MR49F PCa cells subjected to siRNA-mediated knockdown of DECR2 (**Figure 4a**). Inspection of the lipid profiles of LNCaP, V16D and MR49F PCa cells after transient knockdown of DECR2 revealed a profound remodelling of the cellular lipidome (**Figure 4b**). All 3 cell lines displayed a strong and consistent accumulation of many cellular lipids (**Figure 4b; Supplementary Figure 4c; Supplementary Data 2**), in particular the cholesteryl esters, glycerides, sphingolipids, and several classes of (lyso)-phospholipids such as PC, PE, PI and LPE (**Figure 4b**; **Supplementary Figure 4e**). Interestingly, we also observed the accumulation of several classes of ether-linked phospholipids such as PC-O, PE-O and PE-P (**Figure 4b**; **Supplementary Figure 4e**), which are known to be synthesised within the peroxisome^31^. DECR2 knockdown did not markedly alter saturated fatty acids (SFA) levels, but significantly increased the abundance of monounsaturated (MUFA) and polyunsaturated fatty acids (PUFA) compared with control cells (**Figure 4c**). The opposite effect on lipid accumulation was observed in DECR2-overexpressing (hDECR2) LNCaP cells, whereby we observed a significant reduction in total lipid levels (**Figure 4d**; **Supplementary Figure 4d; Supplementary Data 2**). Analysis of the lipidome revealed marked decrease in abundance across almost all classes of lipids compared with control cells, except triacylglycerides (TAG) which showed a significant increase in levels (**Figure 4d**; **Supplementary Figure 4f**). Overexpression of DECR2 in LNCaP cells markedly decreased SFA, MUFA and PUFA abundance compared with control cells (**Figure 4e**). However, closer inspection of the relative abundance revealed a significant decrease in the proportions of SFA and PUFA (n ≥ 3) levels relative to MUFA and PUFA (n = 2) levels (**Figure 4f**). Next, we performed a lipid ontology^32^ (LION) enrichment analysis to associate lipids in DECR2 knockdown cells with biological features (**Figure 4g**). Lipids with increased relative abundance in DECR2 knockdown samples were enriched for terms associated with lipid storage/droplet, sphingolipids, lipid-mediated signalling, PG, PI and PS, and alterations in lipids implicated in the plasma membrane and endosome/lysosome. In contrast, LION enrichment analysis revealed enrichment for terms associated with PUFAs, SM, PC, PE, plasmalogens, and membrane components such as the mitochondria and endoplasmic reticulum, among lipids downregulated in DECR2 knockdown cells.

**Figure 4.**
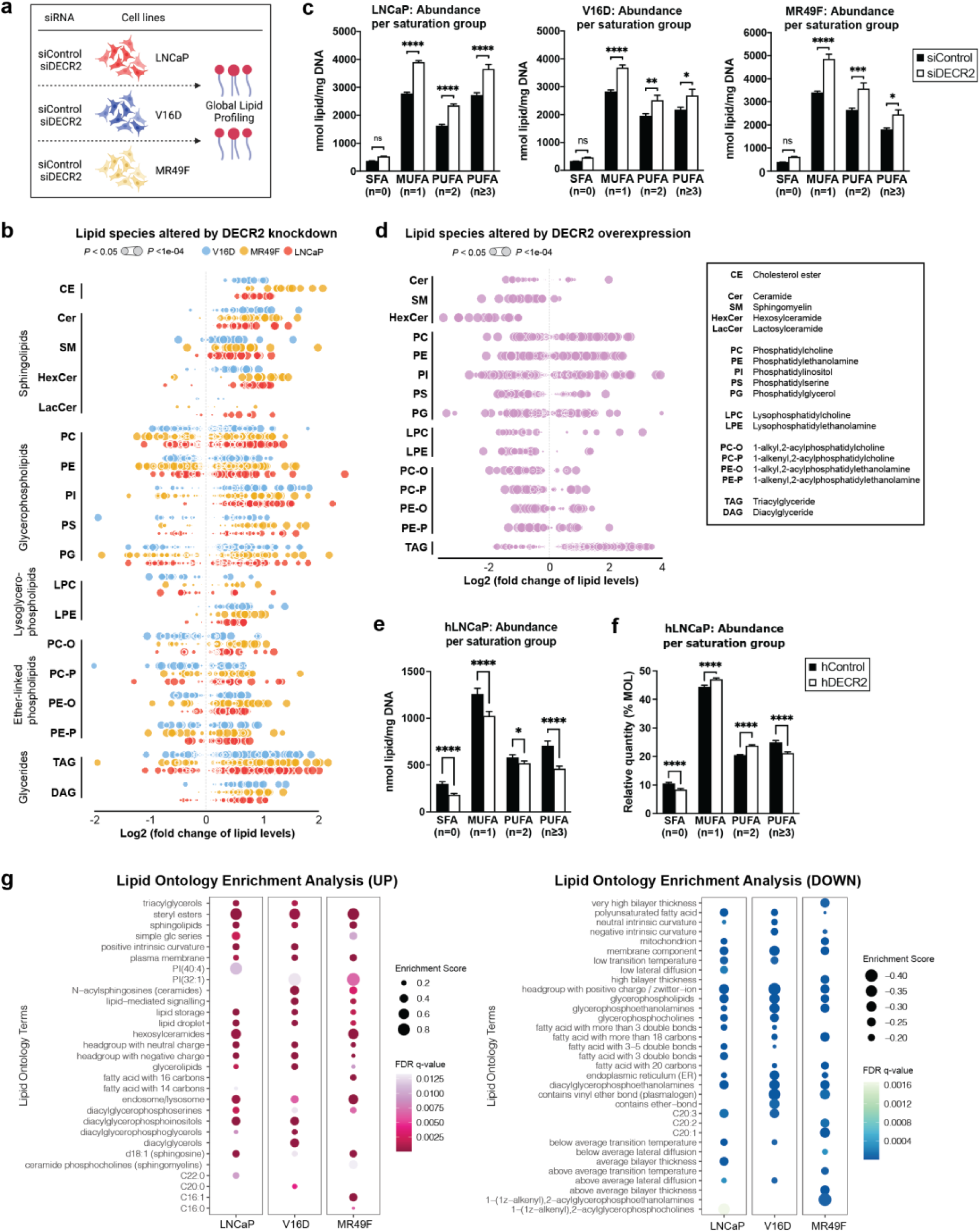
Global lipidomics of DECR2 knockdown and overexpression in prostate cancer cells reveal strongly altered lipid states. **(a)** Global lipidomic study overview. **(b)** Lipidomic analysis of LNCaP, V16D and MR49F prostate cancer cell lines subjected to siRNA-mediated DECR2 knockdown relative to control. Values are shown as quantitative log2-fold changes. Lipidomics data were from 6 replicates and are represented as means. Each dot represents a lipid species. Dot size is proportionate to statistical significance (see **Supplementary Data 2**). **(c)** Quantitative abundance per saturation group in LNCaP, V16D and MR49F cells. **(d)** Lipidomic analysis of DECR2 overexpression (hDECR2) cells relative to control (hControl). **(e)** Quantitative abundance and **(f)** Relative abundance per saturation group in DECR2 overexpression cells. **(g)** Lipid ontology (LION) enrichment analysis of relative lipid abundance in siControl versus siDECR2 LNCaP, V16D and MR49F prostate cancer cells. Statistical analysis was performed using two-tailed student’s t-test: \**p* < 0.05, \*\**p* < 0.01, \*\*\**p* < 0.001 and \*\*\*\**p* < 0.0001.

### DECR2 expression levels affect sensitivity to enzalutamide

A recent proteomics study by Blomme et al. characterised the changes associated with acquired resistance to AR pathway inhibition (ARi)^2^. Here, we independently analysed their proteomics dataset and found that MSigDB Hallmark and KEGG peroxisomal genes were strongly associated with acquired ENZ and apalutamide (APA) resistance (**Figure 5a**; **Supplementary Figure 5a**). Notably, DECR2 protein was robustly upregulated in ARI-resistant cells and organoids (**Figure 5b**). Further, high DECR2 levels were significantly associated with overall survival (*p* = 0.0233; **Figure 5c**) in the SU2C clinical PCa cohort, consisting of patients with metastatic CRPC linked to longitudinal fatal outcomes^33^. Accordingly, we assessed whether DECR2 depletion or perFAO via TDZ could increase sensitivity of CRPC (22Rv1 and V16D) and ENZ-resistant (MR49F) PCa cells to ENZ. Indeed, DECR2 knockdown further attenuated 22Rv1, V16D and MR49F viability (**Figure 5d**) and colony formation compared to DECR2 knockdown or ENZ alone (**Supplementary Figure 5b**). Similarly, we showed that TDZ in combination with ENZ further attenuated 22Rv1, V16D and MR49F viability (**Figure 5e**) and colony formation (**Supplementary Figure 5c**) compared to TDZ or ENZ alone – most notably in the ENZ-resistant MR49F cells, suggesting re-sensitisation to ENZ (**Supplementary Figure 5c**). TDZ in combination with ENZ also significantly decreased 22Rv1, V16D and MR49F growth in 3D spheroids (**Figure 5f**), which better mimics *in vivo* conditions than 2D cell culture^34^, more effectively than TDZ or ENZ alone. To investigate whether high DECR2 levels could confer resistance to ARi, we assessed cell viability of stable overexpression of DECR2 in LNCaP cells under androgen-depleted conditions. Stable overexpression of DECR2 significantly increased viability of LNCaP cells compared with control cells cultured in charcoal-stripped serum medium (**Figure 5g**). Next, we assessed cell viability of stable DECR2 overexpression LNCaP cells under AR inhibition via treatment with ENZ and APA. In both conditions, stable DECR2 overexpression LNCaP cells were significantly more resistant to ARi compared with control cells (**Figure 5h**).

**Figure 5.**
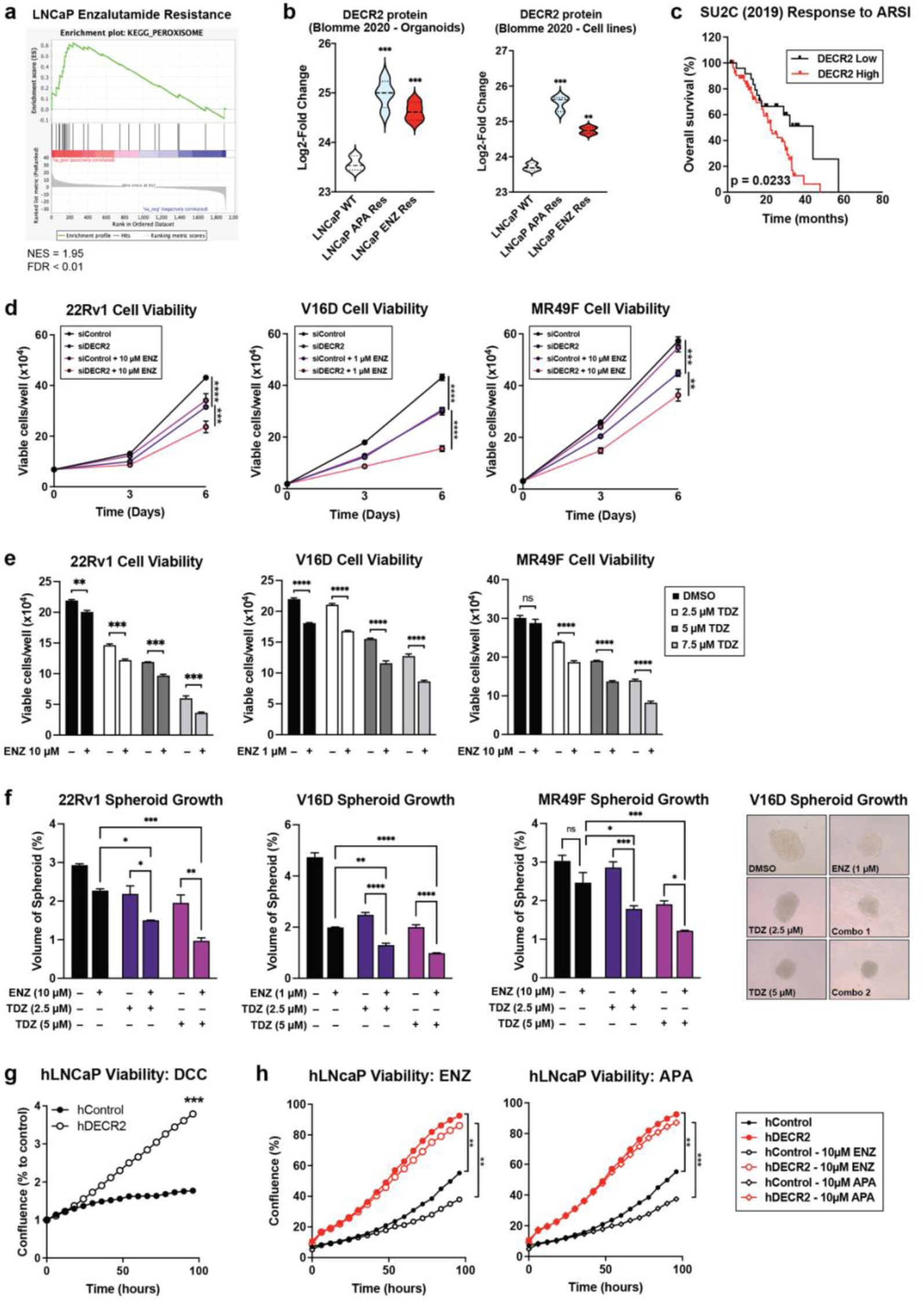
DECR2 confers resistance to enzalutamide in prostate cancer cells. **(a)** GSEA of peroxisomal Hallmark and KEGG proteins shows positive correlation with acquired resistance to enzalutamide. **(b)** DECR2 protein expression is significantly increased in LNCaP acquired apalutamide and enzalutamide resistance organoids and cell lines compared to wildtype LNCaP organoids and cells. Data are represented as violin plots in GraphPad prism. **(c)** The correlation of DECR2 expression with overall survival in the SU2C cohort. ARSI = androgen receptor signalling inhibitor. **(d)** Cell viability of 22Rv1, V16D and MR49F cells subjected to siRNA-mediated DECR2 knockdown, treated with ENZ (1 or 10 µM). **(e)** Cell viability of 22Rv1, V16D and MR49F prostate cancer cell lines treated with thioridazine, TDZ (2.5, 5 and 7.5 µM) and/or ENZ (1 or 10 µM). **(f)** 22Rv1, V16D and MR49F cell growth in 3D spheres, treated with TDZ (2.5 and 5 µM) and/or ENZ (1 or 10 µM). Spheroid volumes were determined after four days of culturing the cells in 96-well microplates; spheres were assessed using the ReViSP software. **(g)** Growth of hDECR2 and hControl LNCaP cells under charcoal-stripped (androgen-deprived) conditions. **(h)** Growth of hDECR2 and hControl LNCaP cells under full serum conditions and in response to enzalutamide (ENZ, 10 µM) and apalutamide (APA, 10 µM). All data are representative of at least 2 independent experiments and presented as mean ± s.e.m of triplicate wells. Data in (c) were statistically analysed using a two-sided log-rank test. Statistical analysis was performed using ordinary one-way or two-way ANOVA. ns = non-significant, \**p* < 0.05, \*\**p* < 0.01, \*\*\**p* < 0.001 and \*\*\*\**p* < 0.0001.

## DISCUSSION

Peroxisomal β-oxidation (perFAO) is an understudied aspect of fatty acid metabolism in PCa. To provide proof-of-principle, and circumvent the time and resources required to develop and test new drug treatments, we appropriated an existing clinically available pharmacological agent to explore the efficacy and clinical exploitability of inhibiting perFAO, using thioridazine (TDZ)^21^. A few studies have examined its anti-tumorigenic effects in several cancer cell types such as the brain^35^, lung^36^, colon^37^, ovarian^38^ and breast^39^. In this study, we provide first-in-field evidence for the therapeutic efficacy of targeting perFAO, using TDZ, in clinical prostate tumours and in *in vitro* cell line models of treatment-resistant PCa. Altogether, our findings suggest that perFAO is an exploitable therapeutic target for PCa. However, recognising that TDZ is not an ideal inhibitor of perFAO due to its lack of specificity and considering that no other perFAO inhibitors currently exist, we sought to identify key functional genes of perFAO that could be more specifically targeted therapeutically.

Cancer-related changes in peroxisomal gene and protein expression and metabolic flux, and their relationship to cellular lipid profile, remain an area ripe for further investigation. Various tumour types exhibit alterations in peroxisome abundance and activity, and it was recently reported that expression of peroxisomal genes is elevated across different tumours^40^. Intriguingly, while some studies observed a decrease in peroxisomal activity in certain tumour types, other groups indicated that peroxisomal metabolic activities promote tumour growth. It is likely that the tumour-promoting or tumour-suppressing functions of peroxisomes are dependent on the tumour type and disease stage^41^. Analysis of the expression of genes involved in peroxisome/β-oxidation in PCa revealed two distinct expression profiles that are either up- or down-regulated with increasing tumour progression.

Not surprisingly, *AMACR* was one of the most consistently upregulated perFAO genes across all stages of PCa, accompanied by *DECR2.* Notably, DECR2 protein expression was highest in castrate-resistant V16D and ENZ-resistant MR49F PCa cell lines. 2,4-dienoyl CoA reductase 2 or DECR2 is a perFAO enzyme analogous to our recently discovered mitochondrial DECR1. Besides residing in different cellular compartments, DECR1 and 2 are both critical NADPH-dependent auxiliary enzymes that play key roles in (poly)unsaturated fatty acid oxidation^42^. Unlike the mitochondria, peroxisomes do not produce ATP. Instead, peroxisomes function to shorten very long chain fatty acids (VLCFA, C≥22) prior to transport into the mitochondria for complete degradation and energy production^31^. Although upon examination of the crystal structure of DECR2 showed selectivity for VLCFAs like docosahexaenoic acid (DHA)^43^, another study reported that DECR2 may be involved in the degradation of short and medium chain substrates^44^. In a very recent study, Spiegel et al. characterised the lipidomic changes of a set of gene knockouts, including DECR2, in a colon cancer cell line. The authors observed elevated levels of long-chain PUFAs in PI, PE-O and PC-O lipid classes (C20 and C22), confirming previous studies that DECR2 can oxidise unsaturated fatty acids and assist in the degradation of PUFAs such as arachidonic acid (AHA) and DHA in peroxisomes^45^.

Our findings shed new light on perFAO in regulating cell cycle progression and lipid metabolism. One of the cellular processes most markedly affected by DECR2 inhibition is the cell cycle, as demonstrated by our RNAseq data. We further showed that DECR2 knockdown markedly arrested the cell cycle at the G1 phase, while high DECR2 levels exhibited an accelerated rate of cell cycle progression. Consistent with our results, we observed a decrease in phosphorylated retinoblastoma (pRb) tumour suppressor protein in DECR2 knockdown compared with control cells. Of particular interest is the connection between peroxisomal fatty acid metabolism and cell cycle progression. We observed that DECR2 knockdown or overexpression had profound effects on the cellular lipidome, notably for altered lipid abundance and composition. Koberlin et al. showed that functional phenotypes could be predicted based on different lipid states, and is applicable to more membrane-dependent processes such as cell division, proliferation, apoptosis, and autophagy^46^. Gokcumen et al. demonstrated that cells actively regulate their lipid composition and localisation during cell division, and that specific lipids within lipid families have specific functions that contribute to signalling and structural integrity of dividing cells^47^. Notably, our LION enrichment analysis revealed dramatic changes in glycerophospholipids in DECR2 knockdown cells, which are essential building blocks for cell growth/proliferation. perFAO contributes to glycerophospholipid (GL) synthesis^48–50^. Previous studies have also shown that key phospholipids of plasma and organelle membrane, PC and PE, are differentially regulated across the cell cycle (G1/S phase) mediated by transcription factors SREBP, AKT/mTOCR1, and p53^51^. perFAO can also alter the GL milieu to trigger actin remodelling to enable the reorganisation of the plasma membrane needed for proper receptor localisation, recruitment of signalling intermediates and changes in cell morphology required for proliferation and survival^52^. Certain lipid species such as DGs and sphingolipids (ie. SM and Cer) can act as second messengers, and changes in the acyl-chain composition of membrane lipids such as PC are known to impact the regulation of oncogenic signalling pathways^53^. Some studies have linked Rb/pRb to the control of lipid metabolism, including but not limited to modulation of mitochondrial oxidative phosphorylation^54^ and lipid remodelling (ie. elongation and desaturation)^55^. In addition, Rb/pRb is able to cooperate with various metabolic pathways (ie. mTORC1, SREBP, PI3K/AKT) to facilitate homeostatic control of cellular metabolism^56^. Intriguingly, we observed a marked increase in TG levels in DECR2 knockdown cells, suggesting an accumulation of lipid droplets (LD) which we confirmed via BODIPY 493/503 neutral lipid staining. Previous studies have shown that LDs are tightly linked to cell cycle progression, particularly in the G1/S phase transition, either by increasing their interaction with the mitochondria and peroxisomes, or to microtubules, to energetically fuel cell proliferation and promote cell survival through increased fatty acid oxidation, or to maintain lipid homeostasis for efficient cell division^57^. More recently, perFAO has been shown to prevent lipotoxicity by regulating lipolysis from LDs via control of ATGL protein levels^58^. Alterations in membrane lipid composition have also been implicated to alter membrane properties such as membrane fluidity, in a way that promotes survival and treatment resistance in cancer cells^59^. This can be attributed to changes in the desaturation ratio of membrane lipids, increased SM and/or cholesterol content, and the formation of detergent-resistant membrane domains that can activate multi-drug efflux transporters^60^. Acyl-chain lengths can also have a profound impact on curvature, fluidity, and fusion rates of biological membranes, although the biological roles of different chain lengths and their regulation are less understood. Some possibilities include remodelling and expanding the endoplasmic reticulum (ER)^61,62^, and mobilisation of lipids from LD stores^51,63,64^, supporting the idea that membrane synthesis and integrity could have potential impacts on cell division. In PCa, it was recently shown that fatty acid chain elongation via ELOVL5 promotes prostate tumour growth^65^. Future work is warranted to explore in detail and dissect the specific roles of DECR2 in these activities and the possibilities that these functional pathways intersect in order to drive tumour progression.

Besides uncovering a fundamental role for DECR2 in regulating lipid homeostasis and cell cycle regulation, this study highlights the importance of perFAO for the first time in CRPC or treatment resistance. Our findings suggest that abnormal perFAO is likely to be one of the contributing factors for resistance to enzalutamide. Herein, we showed that combination of DECR2 inhibition or TDZ with enzalutamide further abrogated PCa cell proliferation of 22Rv1, V16D and MR49F PCa cells and overexpression of DECR2 confers LNCaP cells to be more resistant to enzalutamide. Interestingly, a study by Shen et al. reported that BRAF mutant melanoma ‘persister’ cells resistant to BRAF/MEK inhibition switch their metabolism from glycolysis to oxidative phosphorylation that is predominantly dependent on perFAO compared to mitochondrial β-oxidation^22^. Peroxisomal-derived ether-linked phospholipids have been shown to drive susceptibility to and evasion from ferroptosis^66^. Gajewski et al. found that DECR2 loss promotes the resistance of tumour cells to immunotherapy by evading CD8+ T-cell-mediated tumour ferroptosis *in vivo*^67^. This suggests that low DECR2 expression may be advantageous to promote tumour growth, which is contradictory to our current findings. More extensive work will be needed to understand the diversity of mechanisms involved in the regulation of DECR2 in PCa progression and treatment resistance. It is important to recognise that peroxisomes do not function as an isolated entity. Organelle crosstalk and functional interplay exists between peroxisomes and many organelles such as the mitochondria, ER, lipid droplets and lysosomes to maintain metabolic homeostasis^68^. A major challenge will be to reveal the mechanisms that mediate the metabolic interplay between peroxisomes and other organelles, and how these are impacted in various diseases/disorders. Our study also extends the current focus of peroxisome-mediated lipid changes in cancer cells to exploring the contribution of peroxisomes to tumorigenesis in the tumour microenvironment (i.e. immunity and inflammation^52,69,70^), potentially opening up new therapeutic avenues to fight tumour cell proliferation by targeting peroxisome-related processes. Collectively, our findings make a new contribution to the study of altered lipid metabolism in PCa and reveals DECR2 as a major modulator of cell cycle progression and lipid metabolism, and an exciting novel candidate for therapeutic targeting.

## MATERIALS AND METHODS

### Cell lines and tissue culture

Human immortalized normal prostate epithelial cell lines PNT1 and PNT2 were obtained from the European Collection of Authenticated Cell Cultures (ECACC). Prostate carcinoma cells LNCaP and 22RV1 were obtained from the American Type Culture Collection (ATCC; Rockville, MD, USA). Castrate-resistant V16D and enzalutamide-resistant MR49F cell lines were derived through serial xenograft passage of LNCaP cells^71^ and were a kind gift from Professor Amina Zoubeidi’s laboratory (Vancouver Prostate Centre, Vancouver, Canada). All cell lines were verified in 2022 via short tandem repeat profiling (Cell Bank Australia). Cells were cultured in RPMI-1640 medium containing 10 % fetal bovine serum (FBS; Sigma-Aldrich, NSW, Australia) in a 5 % CO_2_ humidified atmosphere at 37 °C; 10 µM enzalutamide was supplemented in the media for growth of MR49F cells.

### Ex vivo culture of human prostate tumours

Patient derived-explant (PDE) culture was carried out according to techniques established in our laboratory, as described previously^27^. Briefly, 6 mm/8 mm biopsy cores were collected from men undergoing robotic radical prostatectomy at St. Andrew’s Hospital (Adelaide, South Australia) with written informed consent through the Australia Prostate Cancer BioResource. Tissues were dissected into smaller 1 mm^3^ pieces and cultured on Gelfoam sponges (80 x 125 mm Pfizer 1205147) in 24-well plates pre-soaked in 500 μL RPMI-1640 medium supplemented with 10 % FBS and antibiotic/antimycotic solution. TDZ (10 μM or 20 μM) was added into each well and the tissues were cultured in 5 % CO2 in humidified atmosphere at 37 °C for 48 h, then snap frozen in liquid nitrogen and stored at −80 °C, or formalin-fixed and paraffin-embedded. Clinicopathological features of the patients included in this study are shown in **Table 1**.

### Immunohistochemistry (IHC)

Paraffin-embedded tissue sections (2 – 4 µm) were prepared prior to staining as previously described. IHC staining was performed using DECR2 (ab153849 Abcam, diluted 1:1000) antibody and the 3,3’-Diaminobenzidine (DAB) Enhanced Liquid Substrate System tetrahydrochloride (Sigma Aldrich) as described previously. DECR2 staining intensity was measured by Video Image Analysis.

### Analysis of publicly available prostate cancer datasets

Gene expression data were downloaded from The Cancer Genome Atlas (TCGA) data portal, cBioPortal (SU2C and MSKCC)^72^, and the GEO website; Taylor et al, GSE21034^19^; Grasso et al, GSE35988^20^; Tomlin et al, GSE6099^28^. Proteomics data^2^ (raw MaxQuant files) was downloaded from the ProteomeXchange Consortium via the PRIDE partner repository using the dataset identifier PXD016836 and analysed independently using the R software version 3.6.3.

### Western blotting

Protein lysates were collected in RIPA lysis buffer (10 mM Tris, 150 mM NaCl, 1 mM EDTA, 1 % Triton X-100, 10 % protease inhibitor). Western blotting on whole cell protein lysates were performed as previously described^73^. Primary antibodies are shown in **Table 2**.

### Quantitative real-time PCR (qPCR)

Total RNA was extracted from cells using the RNAeasy RNA extraction kit (Qiagen), followed by cDNA synthesis using the iScript cDNA Synthesis kit (Bio-Rad) on the CFX384 Real-Time System (Bio-Rad, NSW, Australia). qPCR was performed in triplicate as previously described^73^. Relative gene expression was determined using the comparative Ct method and normalised to the internal housekeeping genes *GUSB* and *L19*. Primer sequences are shown in **Table 3**.

### siRNA transfection

Human DECR2 ON-TARGET plus SMART pool (L-009627-00-0005) small interfering RNAs (siRNAs) and control siRNA (D-001810-01-20 ON-TARGET plus non-targeting siRNA #1) were purchased from Millennium Science (Victoria, Australia). siRNAs (5 nM) were reverse transfected using Lipofectamine RNAiMAX transfection reagent (Invitrogen, Victoria, Australia) according to manufacturer’s instructions.

### Generation of stable shDECR2 and hDECR2 LNCaP cells

LNCaP cells were transduced with the universal negative control shRNA lentiviral particles (shControl) or hControl (GFP-Puro), DECR2 shRNA inducible lentiviral particles (shDECR2, RFP-Puro) designed by Horizon Discovery (Cambridge, UK), or hDECR2 (GFP-Puro) designed by GenTarget Inc (San Diego, CA, USA) according to manufacturer’s instructions.

### Functional assays

#### Cell viability

Cells were seeded in triplicate in 24-well plates at a density of 2.5 x 10^4^ – 6.0 x 10^4^ cells/well and reverse transfected with siRNA overnight or treated with drug supplemented medium. Cells were manually counted using a hemocytometer 96 h post-siRNA knockdown or treatment and cell viability was assessed by Trypan Blue exclusion as described previously^73^.

#### Cell proliferation / growth

Cells were seeded in 96-well plates at a density of 3 x 10^3^ – 5 x 10^3^ cells/well and treated with drug supplemented medium. Plates were then placed in the IncuCyte® live-cell analysis system (Sartorius) and images of the cells were recorded every 6 hours. Cell growth or proliferation were determined as a measure of confluency using the confluence image mask on the Incucyte® Base Analysis Software.

#### Cell migration

Transwell migration assays were performed using 24-well polycarbonate Transwell inserts (3422, Sigma-Aldrich). C42B and 22RV1 cells transfected overnight with siRNA were seeded into the upper chamber of the Transwell at a density of 9.0 x 10^4^ – 1.5 x 10^5^ cells/well in serum-free medium. 650 μL of medium containing 10 % FBS was added to the bottom chamber. Cells were incubated at 37 °C for 48 h. For TDZ treatment, medium in both the upper and lower chambers were supplemented with TDZ (2.5 μM). Inserts were washed with PBS and non-migrated cells were gently removed using a cotton-tipped swab. The inserts were then fixed in 4 % paraformaldehyde for 20 min and stained with 1 % crystal violet for 30 min. Images of migrated cells were captured using the Axio Scope A1 Fluorescent Microscope (Zeiss) at 40 X magnification. The number of migrated cells were counted manually and presented as percentages relative to control cells ± SEM.

#### Colony formation assay

DECR2 stable knockdown (shDECR2) cells or DECR2 overexpression (hDECR2) cells were prepared in a single-cell suspension before seeding in 6-well plates at a density of 500 cells/well. For TDZ treatment, C42B, V16D and MR49F cells were seeded overnight and gently treated with drug-supplemented medium. Cells were incubated for 2 weeks at 37 °C with medium being replenished every 3-5 days. After 2 weeks, cells were washed with PBS and fixed with 4 % paraformaldehyde, then stained with 1 % crystal violet for 30 min. Colonies were counted manually and results were reported as number of colonies ± SEM.

#### 3D Spheroid growth assay

For TDZ treatment, 22Rv1, V16D and MR49F cells were seeded (400 – 700 cells/well) overnight in Nunclon Sphera 96-well U-shaped-bottom microplates (Thermo Fisher) and gently treated with drug-supplemented medium. Cells were incubated for 6 days at 37 °C. Images of spheroids were captured, and the sphere volume was determined using ImageJ and the ReViSP software^74^.

### Flow cytometry

#### Cell cycle analysis

Cells were seeded in triplicate in 6-well plates at a density of 3 x 10^5^ – 6 x 10^5^ cells/well and reverse transfected with siRNA overnight or treated with drug supplemented medium. Cells were collected into fluorescence-activated cell sorting (FACS) tubes and centrifuged at 1,500 rpm for 5 min, then fixed in cold 70 % ethanol for 2 h. Samples were stained with 50 μg/mL of propidium iodide (PI, Sigma-Aldrich) and 100 μg/mL Ribonuclease A from bovine pancreas (Sigma-Aldrich) for 30 min at room temperature. Cells were analysed using a BD LSRFortessa X-20 Flow Cytometer (BD Biosciences). Data were evaluated using FlowJo version 10.

#### Apoptosis assay

Cells were seeded in triplicate in 6-well plates at a density of 3 x 10^5^ – 6 x 10^5^ cells/well and reverse transfected with siRNA overnight or treated with drug supplemented medium. Cells were collected into FACS tubes and centrifuged at 1,500 rpm for 5 min, then resuspended in FACS Binding Buffer (94 % Hank’s Balanced Salt Solution, 1 % HEPES, 5 % CaCl_2_), 7-AAD (Thermo Fisher Scientific) and Annexin-V PE (BD) for 30 min in the dark. Cells were analysed using a BD LSRFortessa X-20 Flow Cytometer (BD Biosciences). Data were evaluated using FlowJo version 10.

#### Neutral lipid content quantification

Cells were seeded in triplicate in 24-well plates at a density of 3 x 10^5^ – 6 x 10^5^ cells/well and reverse transfected with siRNA overnight or treated with drug supplemented medium. Cells were collected into FACS tubes and centrifuged at 1,500 rpm for 5 min, then resuspended in 2 μM of fluorescent neutral lipid dye BODIPY 493/503 (Thermo Fisher Scientific) for 15 min at 37 °C. Cells were resuspended in 300 μL FACS Binding Buffer and analysed using a BD LSRFortessa X-20 Flow Cytometer (BD Biosciences). Data were evaluated using FlowJo version 10.

#### Lipidomics

Lipid extraction, mass spectrometry, and data analysis methods were performed as previously described^12^. Unpaired t-test p-values and FDR corrected p-values (using the Benjamini/Hochberg procedure) were calculated using R version 4.1.2 and visualised using ggplot2 version 3.3.5.

### RNAseq

#### RNA extraction and library preparation

For RNAseq, 6 biological replicates of V16D and MR49F prostate cancer cells subjected to either siControl or siDECR2 knockdown for 72 h were analysed. Total RNA was extracted using TRIzol reagent (Thermo Fisher) and the RNeasy Micro Kit (Qiagen) and then depleted for DNA using RNase-Free DNase Set (Qiagen). RNA quality and quantity was determined using the Tapestation 2200 and Qubit, respectively. Libraries were generated using the Nugen Universal Plus mRNA-seq protocol and converted to MGI compatible libraries using the MGIEasy Universal Library Conversion Kit. Libraries were sequenced on the MGI DNBSEQ G400 (paired-end reads, 2 x 98 bp) at the South Australian Genomics Centre (SAGC), South Australian Health and Medical Research Institute, Australia.

#### RNAseq analysis

Sequence read quality was assessed using FastQC version 0.11.3 and trimmed with Trimmomatic version 0.36 with a sliding window of a minimum PHRED score of 20 and a window size of 4 nucleotides. Reads were also filtered for a minimum length of 36 nucleotides. Next, reads were aligned to GRCh38 human genome with Ensembl version 105 annotation using STAR version 2.7.9a. Gene count matrix was generated with FeatureCounts version subread-2.0.3. Count matrix were imported into R version 4.1.2 for further analysis and visualisation using ggplot2 version 3.3.5. Counts were normalised using the trimmed mean of M values (TMM) method in EdgeR version 3.36 and represented as counts per million (cpm). Differential gene expression analysis was performed using the glmLRT function in EdgeR. Genes with < 2 cpm in at least 25% of samples were excluded from the differential expression analysis. Gene Set Enrichment Analysis (GSEA) was carried out using the GSEA software version 4.2.2 and the Molecular Signatures Database (MSigDB) to identify Hallmark and GO biological processes/pathways that were differentially regulated in the absence of DECR2. Data from GSEA were visualised using the Enrichment Map plugin^75^ in Cytoscape version 3.9.1 to generate the gene interaction network. The resulting network map was filtered using FDR < 0.01 and curated to remove redundant and uninformative nodes, resulting in a simplified network.

### In vivo studies

#### Orthotopic tumour growth (shDECR2)

10 µL containing 1 x 10^6^ DECR2 inducible knockdown cells (shDECR2) were injected intraprostatically in 8-week-old NOD scid gamma (NSG) male mice. Whole-body imaging to monitor luciferase-expressing LNCaP cells was performed at day 3 of the injection and once weekly after that using the In Vivo Imaging System (IVIS, PerkinElmer). Following 1-week post-injection, mice were randomised into two groups: Group A (shControl) mice were fed with sucrose-containing water (2 mg/mL); Group B (shDECR2) mice were fed with doxycycline/sucrose-treated water (2 mg/mL). D-luciferin (potassium salt, PerkinElmer) was dissolved in sterile deionized water (0.03 g/mL) and injected subcutaneously (3 mg/20 g of mouse body weight) before imaging. Bioluminescence is reported as the sum of detected photons per second from a constant region of interest. After the animals were sacrificed, lungs and livers were excised for ex vivo imaging using the IVIS system.

#### Orthotopic tumour growth (hDECR2)

10 μL containing 1 x 10^6^ DECR2 overexpression cells (hDECR2) or negative control cells (hControl) were injected intraprostatically in 8-week-old NSG male mice. Whole-body imaging to monitor luciferase-expressing LNCaP cells was performed at day 3 of the injection and once weekly after that using the In Vivo Imaging System (IVIS, PerkinElmer). D-luciferin (potassium salt, PerkinElmer) was dissolved in sterile deionized water (0.03 g/mL) and injected subcutaneously (3 mg/20 g of mouse body weight) before imaging. Bioluminescence is reported as the sum of detected photons per second from a constant region of interest. After the animals were sacrificed, lungs and livers were excised for ex vivo imaging using the IVIS system.

After each study, tumours that were excised were snap frozen for RNA extraction and formalin fixed and paraffin embedded. All animal procedures were carried out in accordance with the guidelines of the National Health and Medical Research Council of Australia. The orthotopic xenograft studies were approved by the University of Adelaide Animal Ethics Committee (approval number M-2019-037).

### Statistical analysis

Results are reported as mean ± SEM. Statistical analysis was performed using GraphPad Prism (V9.0 for Mac). The differences between treatment groups were compared by t-test or one-way ANOVA followed by Tukey or Dunnett post hoc test, unless otherwise stated in the figure legends. Significance is expressed as \**p* < 0.05, \*\**p* < 0.01, \*\*\**p* < 0.001, \*\*\*\**p* < 0.0001.

## ACKNOWLEDGEMENTS

The results published here are in part based on data generated by The Cancer Genome Atlas, established by the National Cancer Institute and the National Human Genome Research Institute, and we are grateful to the specimen donors and relevant research groups associated with this project. Tissues for the patient-derived explants used in the study were collected with informed consent via the Australian Prostate Cancer BioResource and we thank the doctors, patients and health care professionals involved. We acknowledge expert technical assistance in the study from Samira Khabbazi. Flow cytometry analysis was performed at the South Australian Health Medical Research Institute (SAHMRI) in the ACRF Cellular Imaging and Cytometry Core Facility, generously supported by the Australian Cancer Research Foundation, Detmold Hoopman Group and Australian Government through the Zero Childhood Cancer Program. The authors acknowledge the South Australian Genomics Centre (SAGC) which provided the RNA-sequencing services. The SAGC is supported by the National Collaborative Research Infrastructure Strategy (NCRIS) via BioPlatforms Australia and by the SAGC partner institutes. Animal studies were performed at the Bioresources Facilities at the South Australian Health and Medical Research Institute. The authors also thank Adelaide Microscopy (University of Adelaide).

## AUTHOR CONTRIBUTIONS

CYM, ZDN and LMB conceived the study and wrote the manuscript. CYM, MH, DJL, JD, JVS designed experiments. CYM, ADTN, TN, JD, NR, JVS acquired data and performed experiments. CYM, ADTN, TN, FR, DJL interpreted and analysed the data. DJL, ZDN, LMB supervised the study. JVS, ZDN, LMB acquired funding. MH, CYM performed the *in vivo* experiments. All authors read the manuscript, agree with the content, and were given the opportunity to provide input.

---

**Supplementary Figure 1.**
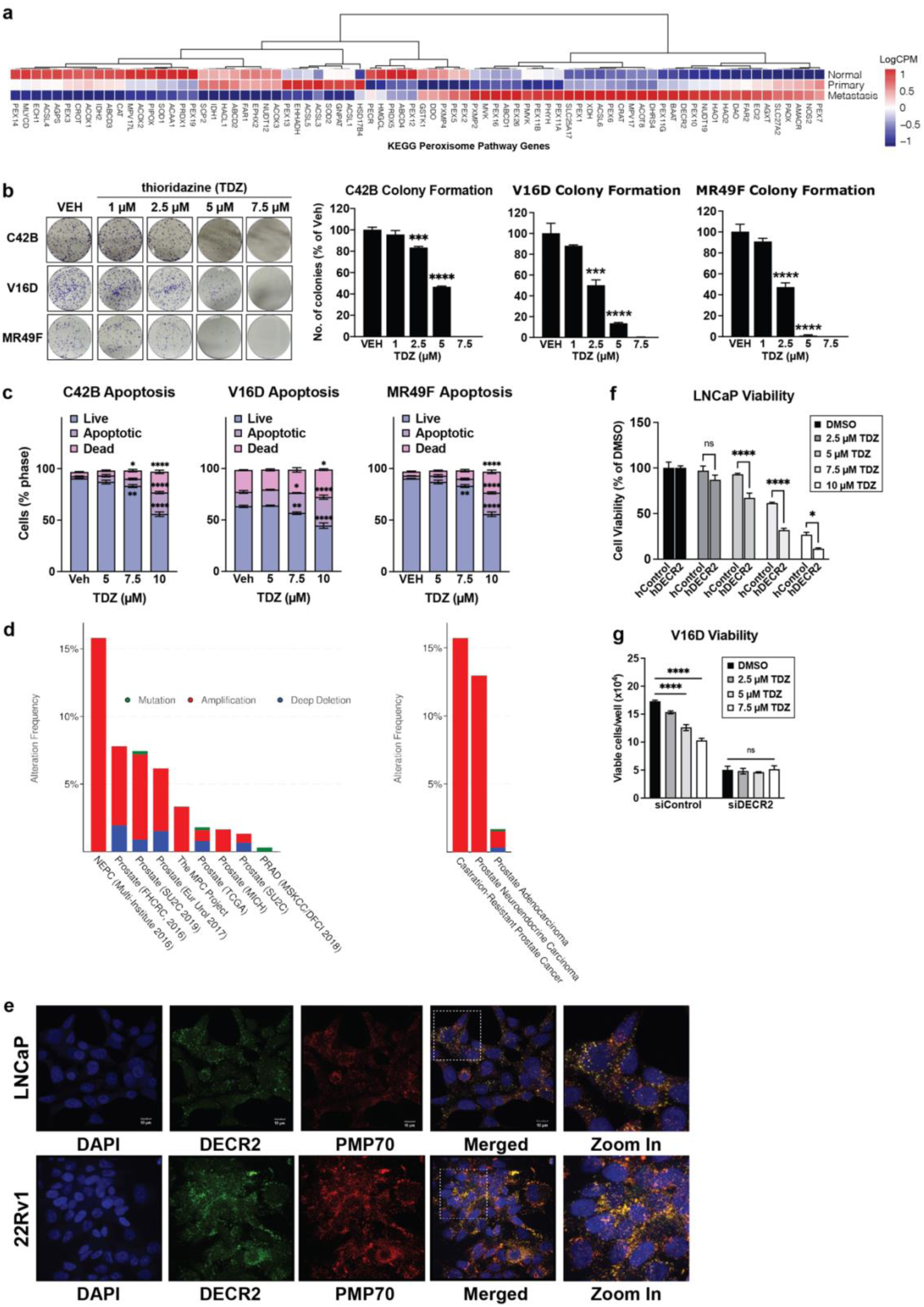
**(a)** Heatmap of KEGG peroxisome pathway genes in the Taylor cohort. **(b)** C42B and 22Rv1 prostate cancer cell lines treated with TDZ (2.5 μM) were assessed for cell migration using a transwell migration assay. **(c)** C42B, V16D and MR49F prostate cancer cells were treated with TDZ for 48 h and assessed for apoptotic and dead cells via flow cytometry. Data presented as percentage of cells in each live, apoptotic, or dead state per sample. **(d)** Histogram displaying DECR2 mutation and copy-number alteration frequency across 9 prostate cancer genomic datasets (left), and across 3 prostate cancer subtypes (right). **(e)** Immunocytochemistry staining of LNCaP and 22Rv1 cells to determine subcellular localisation of DECR2. DAPI: nuclei; Alexa Fluor 488 secondary antibody: DECR2; Alexa Fluor 594 secondary antibody: PMP70 (Peroxisome), scale bar = 10 µm. **(f)** Cell viability of overexpression hDECR2 cells versus hControl LNCaP cells after treatment with varying doses of TDZ. **(g)** Viability of V16D cells subjected to siRNA-mediated DECR2 knockdown with or without TDZ treatment. All cell line data are representative of at least 2 independent experiments and presented as mean ± s.e.m of triplicate wells. Statistical analyses were performed using ordinary one-way or two-way ANOVA: \**p* < 0.05, \*\**p* < 0.01, \*\*\**p* < 0.001 and \*\*\*\**p* < 0.0001.

**Supplementary Figure 2.**
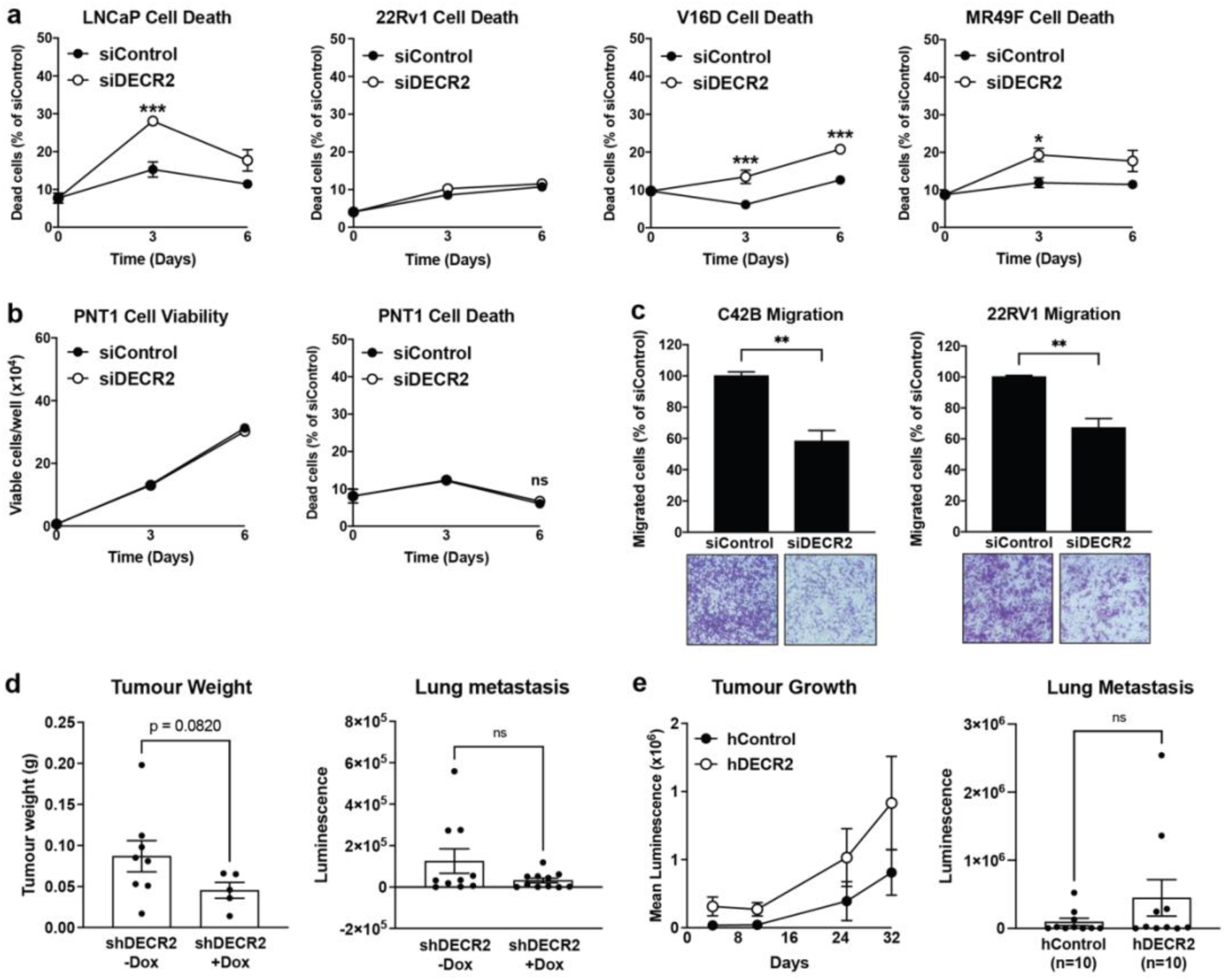
**(a)** Cell death of androgen-dependent LNCaP, castrate-resistant 22Rv1 and V16D, and enzalutamide-resistant MR49F prostate cancer cell lines subjected to siRNA-mediated DECR2 knockdown. **(b)** Cell viability and cell death of non-malignant prostate PNT1 cells. **(c)** C42B and 22Rv1 prostate cancer cell lines subjected to siRNA-mediated DECR2 knockdown were assessed for cell migration using a transwell migration assay. **(d)** Tumour weight and lung luminescence readings following DECR2 knockdown in mice (shDECR2 *n* = 11, shControl *n* = 10). Tumour weight includes data from mice with sufficient sized tumours for analysis (shDECR2 *n* = 5, shControl *n* = 8). **(e)** Tumour growth and lung luminescence readings of DECR2 overexpression mice (*n* = 10 per group, including mice with non-detectable tumours). All cell line data are representative of at least 2 independent experiments and presented as mean ± s.e.m of triplicate wells. Statistical analyses were performed using ordinary two-way ANOVA, or two-tailed student’s t-test: ns = non-significant, \**p* < 0.05, \*\**p* < 0.01, \*\*\**p* < 0.001.

**Supplementary Figure 3.**
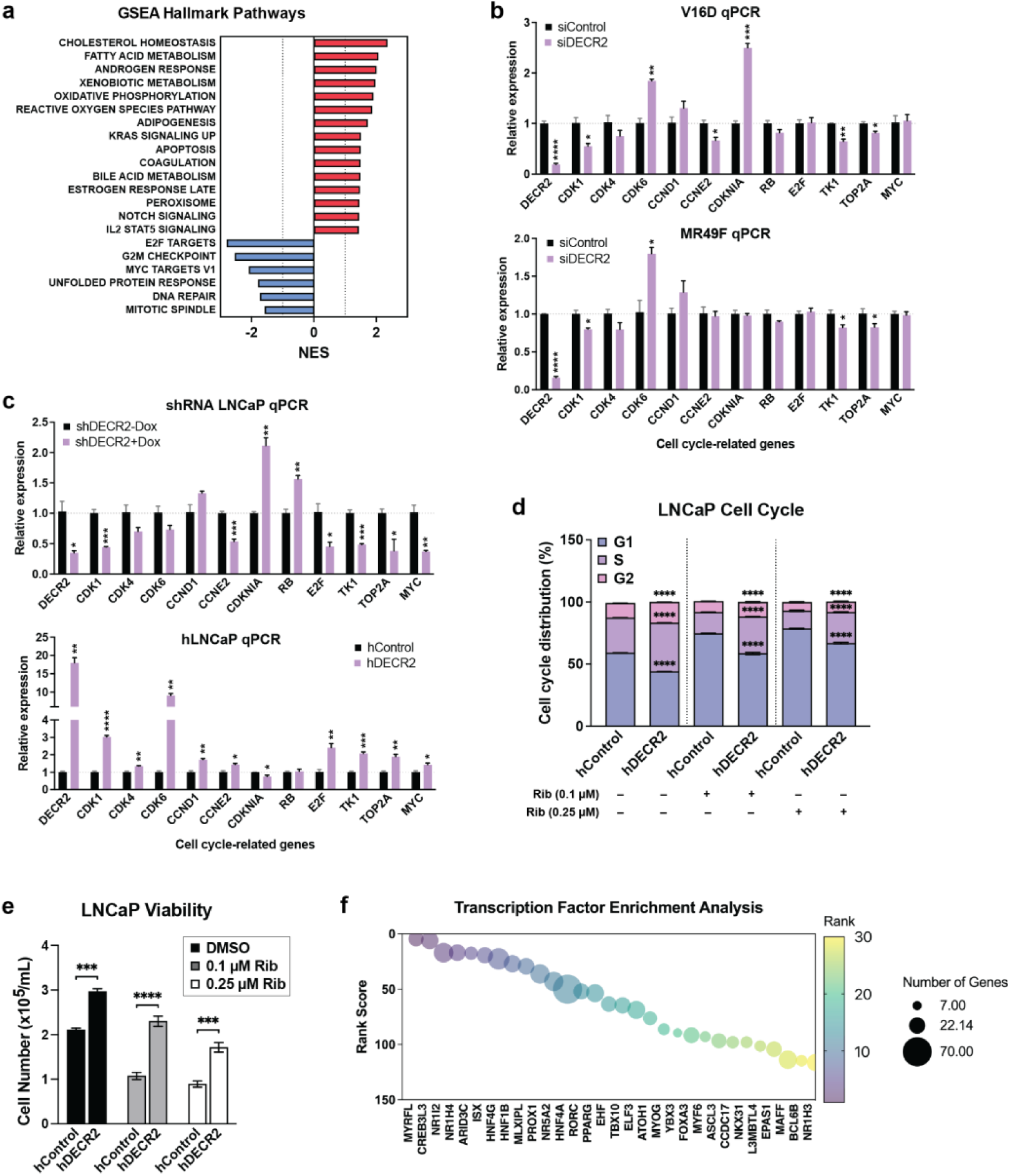
**(a)** Significantly enriched MSigDB Hallmark terms among differentially expressed genes. Quantitative PCR (qPCR) of cell cycle-related genes in **(b)** DECR2 knockdown V16D and MR49F cells, and **(c)** dox-inducible shDECR2 knockdown cells and LNCaP overexpression hDECR2 cells. **(d)** Cell cycle distribution of LNCaP cells with stable overexpression of DECR2, treated with ribociclib (Rib; 0.1 µM and 0.25 µM). **(e)** Viability of LNCaP cells with stable overexpression of DECR2, treated with ribociclib. **(f)** Top 30 transcription factors (TFs) that were enriched in our list of top upregulated differentially expressed genes (*p* < 0.01, log2 fold-change ≥ 1) using the MeanRank method in ChEA3^30^. TFs are ranked from 1 to 30 in ascending order (from left to right), bubble size indicates the number of genes corresponding to the TF targets. All *in vitro* data are representative of at least 2 independent experiments and presented as mean ± s.e.m of triplicate wells. Statistical analyses were performed using ordinary one-way or two-way ANOVA. \**p* < 0.05, \*\**p* < 0.01, \*\*\**p* < 0.001 and \*\*\*\**p* < 0.0001.

**Supplementary Figure 4.**
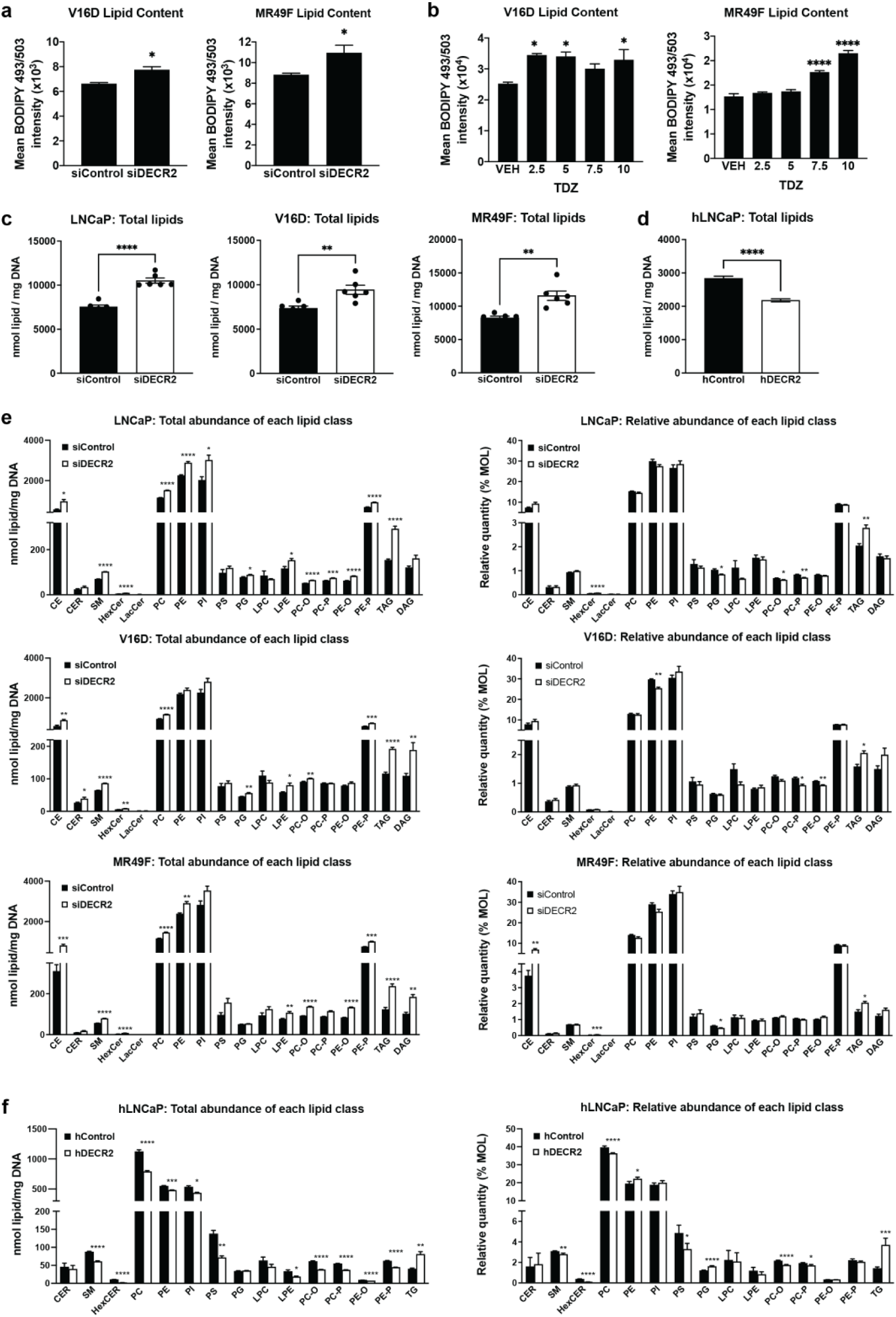
**(a)** V16D and MR49F prostate cancer cells subjected to siRNA-mediated DECR2 knockdown were assessed for neutral lipid content via flow cytometry. **(b)** V16D and MR49F prostate cancer cells were treated with TDZ for 48 h and assessed for neutral lipid content via flow cytometry. Total lipid abundance in **(c)** LNCaP, V16D and MR49F cells, and in **(d)** DECR2 overexpressing LNCaP cells. **(e)** Quantitative (left panel) and relative (right panel) abundance of each lipid class in LNCaP, V16D and MR49F cells. **(f)** Quantitative (left) and relative (right) abundance of each lipid class in DECR2 overexpressing LNCaP cells. Statistical analyses were performed using ordinary two-way ANOVA, or two-tailed student’s t-test. \**p* < 0.05, \*\**p* < 0.01, \*\*\**p* < 0.001 and \*\*\*\**p* < 0.0001.

**Supplementary Figure 5.**
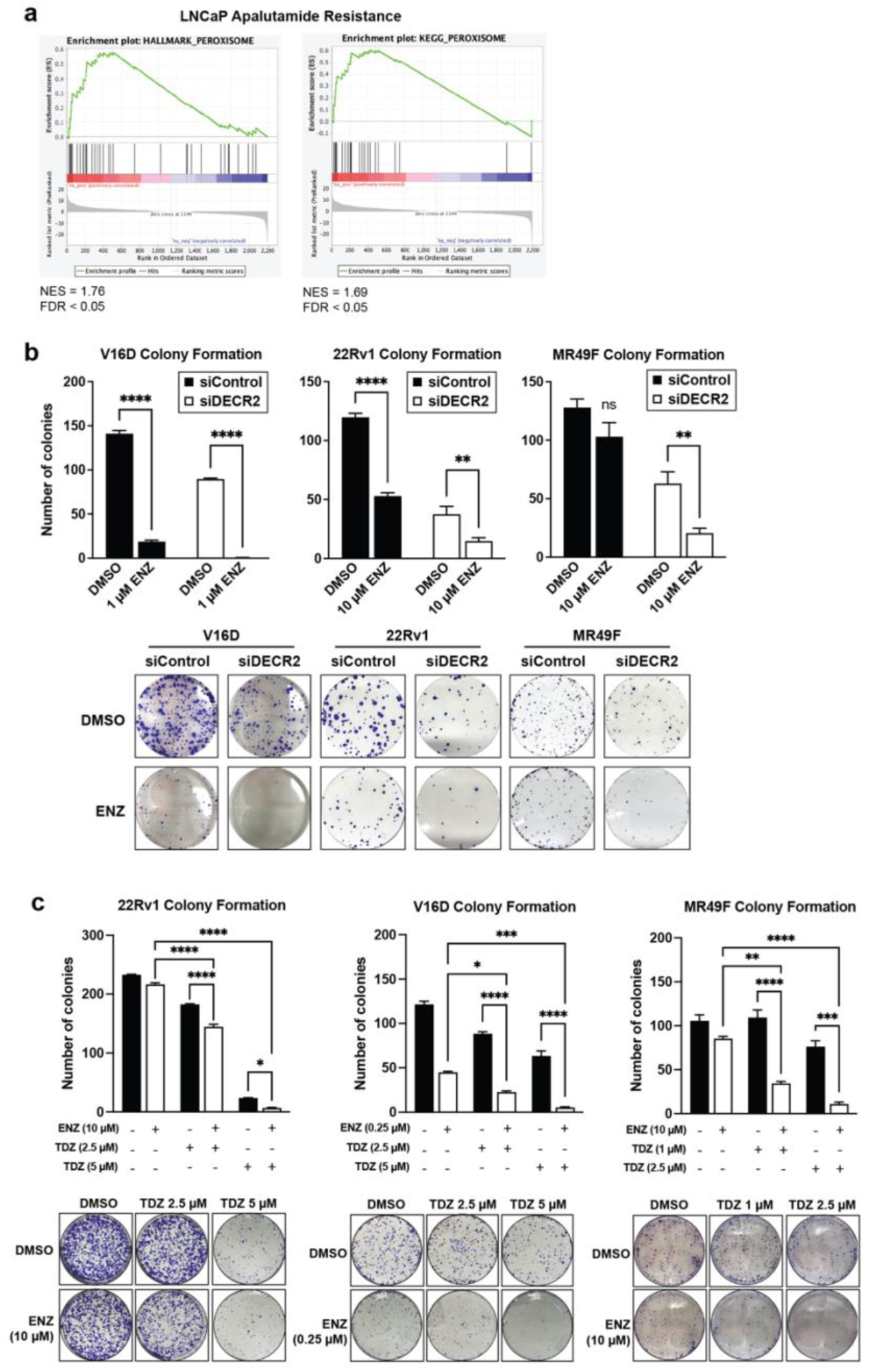
**(a)** GSEA of peroxisomal Hallmark and KEGG proteins shows positive correlation with acquired resistance to apalutamide. **(b)** V16D and MR49F colony formation was evaluated in cells subjected to siRNA-mediated DECR2 knockdown, with or without enzalutamide, ENZ (1 or 10 µM) treatment. **(c)** 22Rv1 and MR49F colony formation was evaluated in cells treated with TDZ (1 and 2.5 µM) and/or ENZ (10 µM). Statistical analysis was performed using ordinary two-way ANOVA. \**p* < 0.05, \*\**p* < 0.01, \*\*\**p* < 0.001 and \*\*\*\**p* < 0.0001.

